# Capturing the Differences between Humoral Immunity in the Normal and Tumor Environments from Repertoire-Seq of B-Cell Receptors Using Supervised Machine Learning

**DOI:** 10.1101/187120

**Authors:** Hiroki Konishi, Daisuke Komura, Hiroto Katoh, Shinichiro Atsumi, Hirotomo Koda, Asami Yamamoto, Yasuyuki Seto, Fukayama Massashi, Rui Yamaguchi, Seiya Imoto, Shumpei Ishikawa

## Abstract

The recent success of immunotherapy in treating tumors has attracted increasing interest in research related to the adaptive immune system in the tumor microenvironment. Recent advances in next-generation sequencing technology enabled the sequencing of whole T-cell receptors (TCRs) and B-cell receptors (BCRs)/immunoglobulins (Igs) in the tumor microenvironment. Since BCRs/Igs in tumor tissues have high affinities for tumor-specific antigens, the patterns of their amino acid sequences and other sequence-independent features such as the number of somatic hypermutations (SHMs) may differ between the normal and tumor microenvironments. However, given the high diversity of BCRs/Igs and the rarity of recurrent sequences among individuals, it is far more difficult to capture such differences in BCR/Ig sequences than in TCR sequences. The aim of this study was to explore the possibility of discriminating BCRs/Igs in tumor and in normal tissues, by capturing these differences using supervised machine learning methods applied to RNA sequences of BCRs/Igs.

RNA sequences of BCRs/Igs were obtained from matched normal and tumor specimens from 90 gastric cancer patients. BCR/Ig-features obtained in Rep-Seq were used to classify individual BCR/Ig sequences into normal or tumor classes. Different machine learning models using various features were constructed as well as gradient boosting machine (GBM) classifier combining these models. The results demonstrated that BCR/Ig sequences between normal and tumor microenvironments exhibit their differences. Next, by using a GBM trained to classify individual BCR/Ig sequences, we tried to classify sets of BCR/Ig sequences into normal or tumor classes. As a result, an area under the curve (AUC) value of 0.826 was achieved, suggesting that BCR/Ig repertoires have distinct sequence-level features in normal and tumor tissues.

To the best of our knowledge, this is the first study to show that BCR/Ig sequences derived from tumor and normal tissues have globally distinct patterns, and that these tissues can be effectively differentiated using BCR/Ig repertoires.

## 1 Introduction

Recent insights into cancer immunity have provided new possible treatment strategies against tumors based on immunotherapy. Since tumor cells contain certain proteins known as tumor-specific antigens (TSAs), which have unique sequences due to somatic mutations and are expressed almost exclusively in tumor environment, evaluation of antigen receptors against TSAs expressed in tumor-infiltrating lymphocytes is important for elucidating cancer immunity. There are two main types of immunity conferred by lymphocytes: cellular immunity, which is largely attributed to the action of T-cell receptors (TCRs), and humoral immunity, which is attributed to the action of immunoglobulins secreted by B-cells.

Recently, advances in next-generation sequencing technology have provided the opportunity to sequence TCRs and B-cell receptors (BCRs) or immunoglobulins (Igs) on an unprecedented scale^1, 2^. However, previous studies analyzing the global patterns of antigen receptor sequences have mostly focused on TCRs^3–7^, resulting in the identification of specific features of the amino acid motifs in TCRs. In contrast, characterizing humoral immunity is more complex than characterizing cellular immunity, because BCRs/Igs show higher sequence diversity than TCRs due to somatic hypermutations. Therefore, analyses of BCRs/Igs have mostly focused on only a small number of known antigens or epitopes^8^. Moreover, the conventional approach used for TCR analysis based on analyzing sequence motifs or identical sequences cannot be applied to BCRs/Igs in tumors, because there are very few BCR/Ig sequences shared by different individuals in cancer microenvironments, unlike infection, vaccine administration, and autoimmunity^9–12^.

Nevertheless, given the importance of humoral immunity in cancer, global sequence analysis of BCRs/Igs in tumors is essential to understand tumor immunity^13^.

We hypothesized that because of TSAs, BCR/Ig sequences in the tumor environment may exhibit characteristics which differ from those in normal tissue environment.

In this study, we tackled this problem by constructing classifiers of BCRs/Igs obtained from the immune repertoire sequencing (Rep-Seq) data of 89-paired tissue specimens obtained from patients with gastric cancer, one of the most common malignancies worldwide and particularly in Asian countries. These classifiers were based on supervised machine learning techniques that differentiated between individual BCR/Ig sequences in normal and tumor environments. V/J-frame pattern, CDR-lengths, the number of SHMs, and physicochemical properties of amino acid sequences of CDRs were used as the key features of BCR/Ig sequences for the differentiation. This approach allowed us to identify distinct characteristics of BCRs/Igs in tumor tissues.

We also classified normal and tumor tissues based on the set of BCR/Ig sequences considering a hypothetical diagnostic situation. Classification of BCR/Ig sequence sets in the context of autoimmune-diseases was conducted previously^14^. Therefore, we compared the classification performance of our classifier with that used in the previous research. This analysis demonstrated that our classifier outperformed the other method when applied to our dataset.

We expect that this approach will advance the field of cancer research and improve immunotherapy toward better personalized medicine in cancer treatment.

## 2 Methods

### 2.1 Clinical samples

Ninety frozen gastric cancer specimens surgically resected from patients between 2009 and 2016 at the University of Tokyo Hospital were analyzed in this study after receiving written, informed consent. This study was approved by the institutional review boards of the University of Tokyo and Tokyo Medical and Dental University.

### 2.2 Rep-Seq data of BCRs/Igs

Total RNA was extracted from approximately 10 sequential frozen sections (10 µm thick) of gastric cancer tissues using the RNeasy Mini kit (Qiagen, Hilden, Germany). Quantity and quality assessments of the extracted total RNAs were made using the Agilent Bioanalyzer (Agilent Technologies, Foster City, CA, USA). Multiplex polymerase chain reaction (PCR) primers targeting BCR/Ig genes (iRepertoire, Inc., AL, USA) were used to amplify BCR/Ig repertoires in each of tumor and normal tissue samples according to the manufacturer’s protocol. Each repertoire library was then sequenced on an Illumina MiSeq instrument (Illumina, San Diego, CA, USA) with 2x 300 bp paired-end sequencing according to the manufacturer’s protocol. The results of paired-end nucleotide sequencing were obtained from two FASTQ files, one read from the 5’ end to the 3’ end and the other from the 3’ end to the 5’ end.

### 2.3 Pre-processing of BCR/Ig sequences

An overview of the workflow for the analysis is provided in Figure 1. The workflow consists of 1) aligning BCR/Ig sequences, 2) defining clonotypes, and 3) extracting dominant BCR/Ig clones.

**Figure 1.**
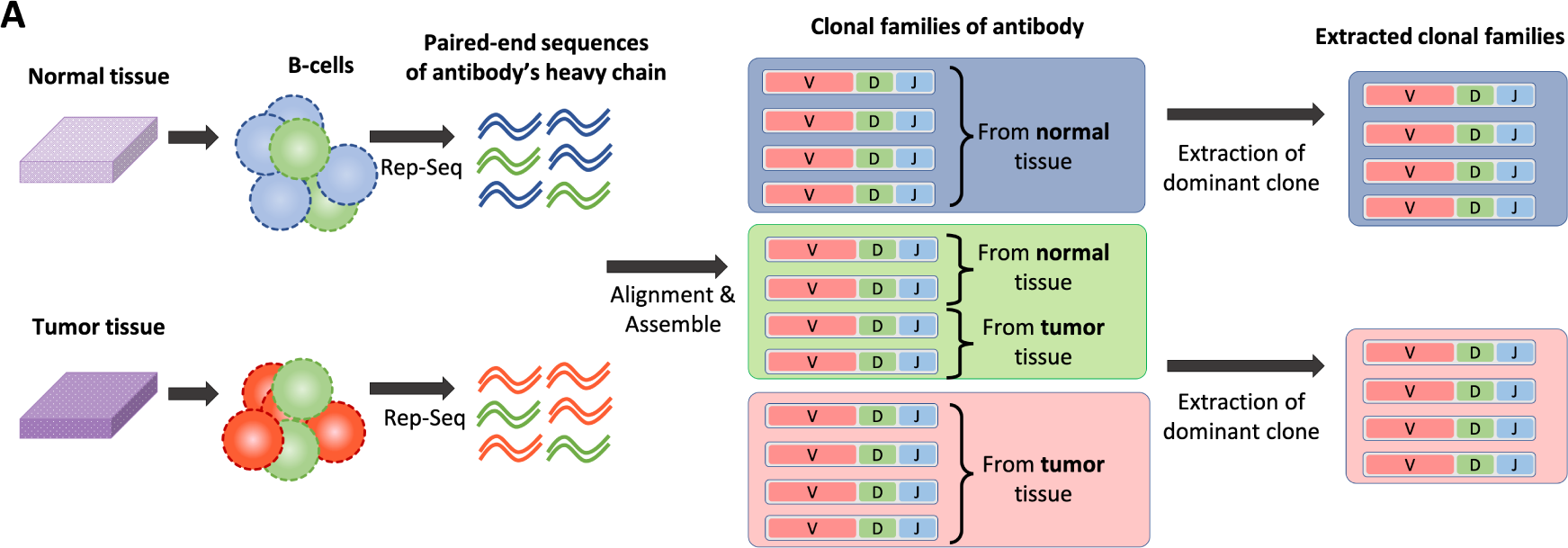
Pipeline for obtaining the BCRs/Igs data used as the query for the classification machine from normal or tumor tissue.

#### 2.3.1 Alignment of BCR/Ig sequences

A heavy chain of BCR/Ig is composed of variable(V), diversity (D), and joining (J) gene segments. To align paired-end nucleotide sequences of BCRs/Igs and estimate the appropriate assignment of V, D, and J gene segments, we used MiXCR, a tool for the universal framework processing of large immunome data from raw sequences for quantitating clonotypes^15^. Since D frame is very short and surrounded by random nucleotides, the accuracy of D frame assignment is generally very low compared to that of V/J frame assignment. Thus, we did not apply D frame assignment to further analysis.

#### 2.3.2 Defining clonotypes

Although the BCR sequence of each B-cell is essentially different, a group of B-cells called a clone has the same ancestral origin, and similar BCR sequences. Clonal lineage of BCRs/Igs was estimated using MiXCR. The following clustering criteria were applied as described by Uduman et al.^16^:

- The same combination of V/J frames
- The same length of CDR3
- A maximum of three nucleotide mismatches in CDR3

BCRs/Igs that satisfy the above criteria were defined as belonging to the same clonal family.

#### 2.3.3 Extraction of dominant BCR/Ig clones

After constructing clonal families, BCR/Ig clones were selected for training the classification machine. Each clone included BCRs/Igs derived from tumor tissues as well as normal tissues. BCR/Ig clones for training were extracted using the following steps: 1) clones with at least 50 reads were extracted to reduce the effect of sequence errors. 2) A clone with 90% or more tumor-derived BCR/Ig content was defined as a tumor-specific clone. The same threshold was applied to define normal clones. 3) Three BCR/Ig sequences were sampled randomly from each extracted clone to eliminate the influence of biased clonal size.

Detailed information about the BCR/Ig data is provided in Table S1

### 2.4 Classification using V/J frame assignment

Classification using V/J frame patterns was conducted using a Bayes classifier. When training the classifier, likelihood of combinations of V/J frames for a given type of tissue (normal or tumor) were calculated as follows

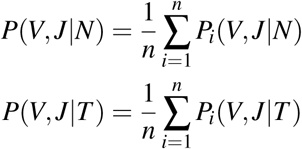

where *n* denotes the number of patients in the training data, and *P*_*i*_(*V, J|N*) and *P*_*i*_(*V, J|T*) denote the relative frequencies of *i*th patient for normal and tumor, respectively. Posterior probabilities of normal or tumor given V/J frames in the test data were calculated using Bayes’ theorem.

Since this classification machine contains no hyperparameter, the performance of the machine is simply measured by conducting leave-one-out cross-validation (LOOCV).

### 2.5 Classification using lengths of CDRs

Classification using lengths of CDRs was conducted in a similar way to the one using V/J frame assignment. Likelihood of combinations of lengths of CDR1, CDR2, CDR3 for a given type of tissue (normal or tumor) were calculated as follows

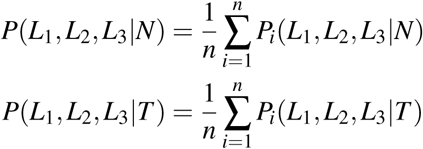

where *L*_1_, *L*_2_ and *L*_3_ denote the length of CDR1, CDR2 and CDR3, respectively, *n* denotes the number of patients in the training data, and *P*_*i*_(*L*_1_, *L*_2_, *L*_3_*|N*) and *P*_*i*_(*L*_1_, *L*_2_, *L*_3_*|T*) denote the relative frequencies of *i*th patient for normal and tumor, respectively. Posterior probabilities of normal or tumor given lengths of CDRs in the test data were calculated using Bayes’ theorem.

### 2.6 Classification using the number of SHMs

Classification using SHMs was conducted using a support vector classifier with linear kernel, and input features are the number of SHM in framework region (FR) and that in CDR.

### 2.7 Classification using amino acid feature vectors

Three kinds of machine learning models were used to classify based on the amino acid sequences of BCRs/Igs: convolutional neural networks (CNN), support vector machine (SVM), and random forest (RF). To take the physicochemical properties of BCRs/Igs into account, we created a feature vector based on Kidera factors, which transforms each amino acid into a 10-dimensional vector. It was originally derived from multivariate analysis of 188 physicochemical properties in each of the 20 amino acids^17^.

Since CDRs in BCRs/Igs are mainly involved in antigen binding, only CDRs are used in this classification. Additionally, their individual lengths were trimmed or padded so that all the feature length be the same. The target lengths of CDRs were determined by their median lengths: 8, 8, and 17 for CDR1, CDR2, and CDR3 respectively. The following methodology was applied to fix the sequence length:

- If the original length was the same as the target length, the sequence was not modified.
- If the original length was larger than the target length, the center of the sequence was trimmed.
- If the original length was smaller than the target length, a pseudo amino acid with a zero feature vector was inserted into the center of the sequence.

The above process created 33(= 8 + 8 + 17)-by-10 matrix for each BCR/Ig. The resulting dimension of the feature vector was 330.

#### 2.7.1 Convolutional Neural Network

The CNN classifier includes a stack of several Conv-Leaky ReLU-MaxPool units, which are followed by fully connected hidden layers with Leaky ReLU activation. The last layer is a fully connected layer with the softmax function for normal and tumor classes. All Leaky ReLU activations were used with leak rate of 0.2. The optimization was done by Adam optimizer with minibatch size set to 100.

Hyperparameters were optimized using a random search strategy^18^. The search range of each hyperparameter is described in Table 1. CNN was implemented using the Python API of TensorFlow^19^.

**Table 1.** Range of hyperparameter searched in the CNN classifier.

| Name of hyperparameter | lower limit | upper limit |
| --- | --- | --- |
| Initial learning rate | 1e-7 | 1e-3 |
| Dropout rate | 0.4 | 1.0 |
| # of convolutional layers | 1 | 2 |
| # of convolutional kernels | 80 | 300 |
| The width of convolution kernels | 2 | 3 |
| The width of pooling filters | 2 | 3 |
| # of fully connected layers | 1 | 3 |
| # of units in a fully connected layer | 100 | 300 |

#### 2.7.2 Support Vector Machine

We used SVM using the radial basis function (RBF) kernel implemented in scikit-learn. There were two hyper-parameters in our SVM classifiers; *C* and *γ*. *C* trades off any misclassification of training examples against the simplicity of the decision surface^20^, and *γ* defines the extent of the influence of a single training example. These hyperparameters were tuned using a grid search strategy. The search range of *C* and *γ* were [10^0^, 10^1^, 10^2^, 10^3^] and [10^−2^, 10^−3^, 10^−4^, 10^−5^,], respectively.

#### 2.7.3 Random Forest

RF implemented in scikit-learn was used^20^. The maximum depth of a tree was tuned as a hyperparameter of the RF model, and its possible values were 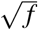, log_2_ *f*, and *f*, where *f* is the number of features (=330) of an input BCR/Ig.

#### 2.7.4 Model selection of machine learning

To optimize the hyperparameters of the classification machines with small number of samples, double cross-validation called nested cross validation was conducted^21^. The purposes of inner and outer cross validation are to determine the hyperparameters and to measure the generalization performance of the determined model, respectively. In our analysis, the inner loop was two-fold cross validation and the outer loop was LOOCV. When holding out validation data in each cross-validation, BCRs/Igs were split at the patient level instead of individual sequence level.

#### 2.7.5 Effect of fixing the length of CDRs

Because the fixed CDR length could cause bias in the classification, effect of CDR length on the performance of our classifier was determined.

To check the effect of trimming and padding the CDR sequences, we calculated the classification performances of each length of CDR3. Because CDR3 has much larger diversity in terms of length and amino acid composition than CDR1 and CDR2, we assumed the effect of trimming and padding would be the largest in CDR3.

### 2.8 Classification using gradient boosting ensemble model

Linear gradient boosting machine implemented in XGBoost was applied to create ensemble classifiers using the four types of features. All hyperparameters were set to default values of Python API of XGBoost.

### 2.9 Motif Analysis

Sequence motifs in CDR1, CDR2, and CDR3 were constructed by applying WebLogo3^22^ to the following 4 groups.

1. BCRs/Igs correctly predicted as Normal tissue-derived with high confidence (*P*(*N*) > 0.9)
2. BCRs/Igs correctly predicted as Tumor-derived with high confidence (*P*(*T*) > 0.9)
3. BCRs/Igs misclassified as Tumor-derived with high confidence (*P*(*T*) > 0.9)
4. BCRs/Igs misclassified as Normal-derived with high confidence (*P*(*N*) > 0.9)

Only sequences with average length of each region (8, 8 and 17 for CDR1, CDR2, and CDR3 respectively) were used.

Both ends of the CDR3 sequence (1^st^ – 3^rd^ amino acids from the N-terminus and 1^st^ – 4^th^ amino acids from the C-terminus), which is highly conserved, were removed to highlight the motifs in the other variable region.

### 2.10 Permutation test

To assess whether or not single BCR/Ig-classifier outperforms random classification, permutation analysis was conducted. The sequence-labels of the BCRs/Igs within each patient were shuffled randomly. A hundred-thousand permutations were performed, and average AUC was calculated over patients in each permutation.

### 2.11 Tissue-level classification

Sets of BCR/Ig sequences were classified into normal or tumor classes.

We used all of the Rep-Seq data of BCRs/Igs when evaluating the classifier in order to simulate actual diagnostic situations, although only tumor/normal-specific clonal families were used when constructing classifiers for individual BCRs/Igs.

Tissue classifier was constructed based on a trained GBM for individual BCR/Ig sequences. Since GBM estimates the probability of individual BCR/Ig deriving from tumor tissue, we used the average probability of the GBM-based classifier for individual BCR/Ig in a set as the tissue-level classification score, which was formulated as follows

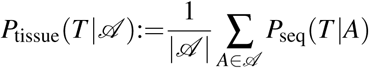

where *A* denotes a vector of outputs of four classifiers using different features for an individual BCR/Ig sequence, and *𝒜* denotes the set of all BCRs/Igs from the same tissue. The classification performance was determined by using a ROC curve on 180 samples from the 90 patients. To classify a target sample from a patient, individual BCR/Ig classifier trained with patients except for the target patient was used. In addition to the average score, tissue-level classification performance was determined by using median or mode of probabilities for comparison.

### 2.12 Training Ostmeyer’s model

Ostmeyer’s model was trained against our dataset from scratch, and evaluated using LOOCV over 90 patients. Training parameters were set to default values except for the number of replicas, which controls the number runs of Adam optimization. The value was set to 1000.

### 2.13 Statistical test

Classification performance (AUC) between classifiers was evaluated using the Wilcoxon signed-rank test for statistical comparisons. These statistical tests were performed using SciPy Python library^23^. *p <* 0.05 was considered significant for all statistical tests. A Bonferroni correction was applied for multiple comparison testing.

To test whether AUCs of different ROC curves differ significantly, the AUCs were compared using R (V.3.4.4) and the pROC package^24^.

## 3 Results

### 3.1 classification of individual BCRs/Igs

First, we investigated whether it is possible to discriminate individual BCR/Ig sequences in normal tissue from those of the tumor tissue.

Table S1 shows the read and clone statistics for each sample we analyzed. Since normal tissues can contain some clones from tumor tissues and vice versa, such contamination in training data could adversely affect the classification performance. Thus, we constructed training data using only tissue type-specific clones for each patient. Clones for 89 patients (1 patient excluded as no tumor-specific clones were identified) were used for the individual BCR/Ig classification. In order to remove the bias of clone sizes, 3 BCR/Ig sequences were extracted from each clone, and as a result, approximately 600,000 BCR/Ig sequences were used as training data in each leave-one-out model. The performance of the classification was measured by AUC-score calculated in each held-out patient in a LOOCV scheme.

We constructed classifiers using BCR features which could contribute to the difference between normal and tumor microenvironment and compared the classification performance among features. The features selected were amino acid sequences of CDRs, lengths of CDRs, V/J-frames, and the number of SHMs. Workflow of the classification using such features is illustrated in Figure 2.

**Figure 2.**
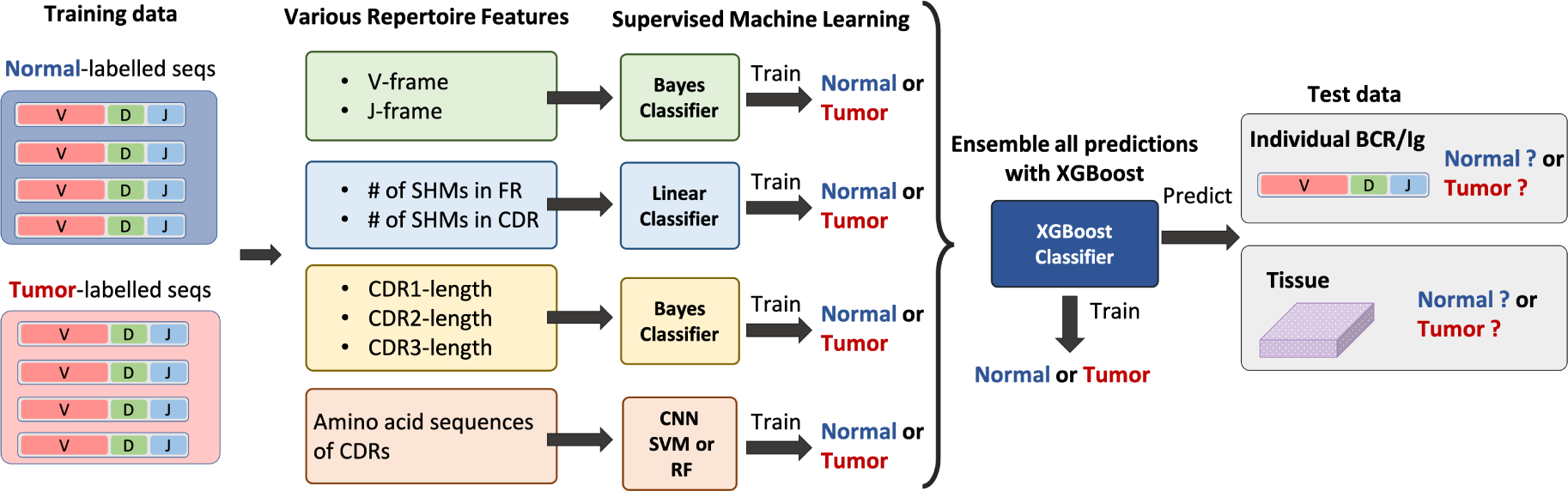
Workflow of individual BCR/Ig classification and tissue classification between normal/tumor environment using various sequence-independent features.

First, multiple classifiers were constructed using amino acid sequences, and their performance was compared. Three representative supervised learning algorithms were compared: Convolutional Neural Networks (CNN), Support Vector Machine (SVM), and Random Forest (RF). To take the physicochemical properties of BCRs/Igs into account, amino acid sequences were encoded into matrices using Kidera factor and used as features. Hyperparameters of these classifiers were tuned using two-fold cross validation. As shown in Figure 3A, CNN significantly outperformed the other two methods. Therefore, CNN using amino acid sequences was adopted in the rest of the study.

In the above experiment, CDR3 length was fixed by trimming or padding amino acid sequences to retain both end of the sequences since there is no straightforward way to deal with variable-length input in CNN, SVM, and RF. To verify our approach, classification performance among different trimming/padding strategies was compared. As shown in Figure 3B, no significant difference was observed between the classification performances of CNN using center-trimmed/padded and ends-trimmed/padded strategies. Figure 3D shows the performance of CNN using each of the various amino acid lengths (8 to 26) of CDR3. Although significant difference was observed (*p* = 0.02, Skillings-Mack test^25^), performances were better than random in all lengths. These results suggest that our trimming/padding strategy does not degrade the performance of CNN.

**Figure 3.**
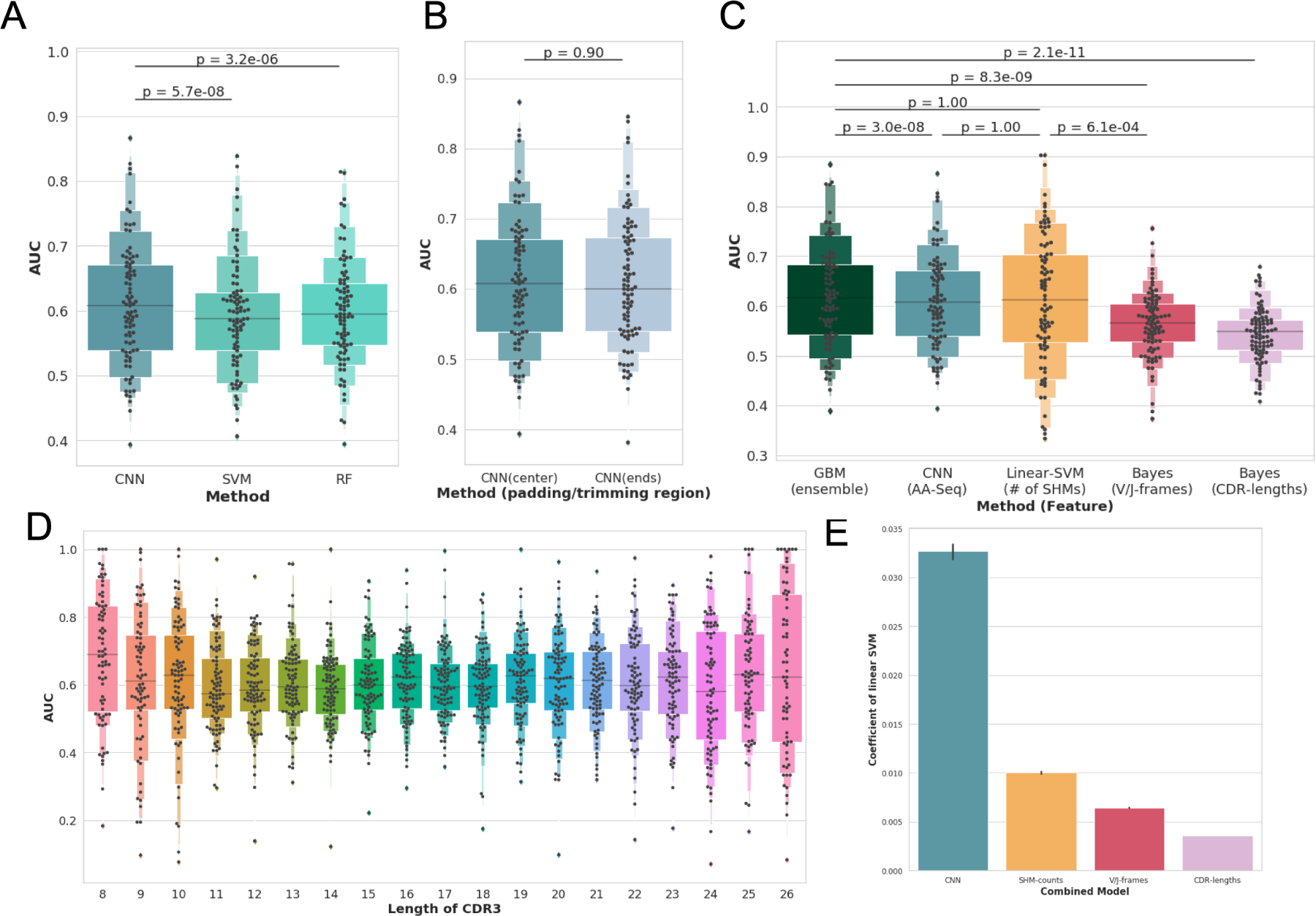
Letter-value plots are showing distribution of the area under the Receiver Operating Characteristic curve (AUROC) calculated on 89 held-out patient. The figures are illustrating the comparison of (A) different models using amino acid sequences, (B) different trimming/padding strategies, (C) models using various sequence-independent features as well as ensemble model combining them, and (D) CNN against different length of CDR3. Barplots (E) shows the coefficients of linear ensemble model.

Our results indicate that BCR/Ig-sequences from normal and tumor environment have some distinct patterns in their amino acid sequences. However, differences in amino acid sequences in normal and tumor environment detected by CNN could be captured by other sequence-independent information, which can be calculated more easily. For example, most of the signals detected by the model may reflect the fact that tumor-derived clones tend to have undergone affinity maturation, while normal tissue-derived clones do not have such tendency. Therefore, classification experiments were conducted using the three sequence-independent features, CDR-length, usage of V/J frames and the number of SHMs, that might also exhibit distinct patterns in normal and cancer tissues.

In order to explore the possibility to improve the classification performance by ensemble, and to evaluate the contribution of each model to the performance, ensemble models were also constructed using multiple features including amino acid sequences by combining all the output probabilities against the sequence from the models described above. Gradient Boosting Machine (GBM) with linear model was used for the ensemble classifier.

Figure 3C shows the performances of different models using different sequence-independent features, as well as the ensemble model. All models performed significantly better than random (p-value < 10^−5^, permutation test), which means that all sequence-independent features we selected have distinct characteristics in normal and tumor microenvironment. Among the four non-ensemble models using sequence-independent features, those using the physicochemical properties of amino acid sequences and number of SHMs showed comparable performances, outperforming the other two models. Although ensemble models outperformed them, the difference was not remarkable. Coefficients of the GBM linear model shown in Figure 3E indicated that output probability of CNN model is the most useful feature for the classification. Nonetheless, the performance of ensemble models that combine all models for each sequence-independent feature was similar to that of CNN, and 89 AUCs of CNN were correlated to those of other classifiers (Figure 4). These results suggest that CNN can deal with the richest information and produce the most reliable probabilities, and that most of the patterns it recognizes might be correlated to other sequence-independent features.

**Figure 4.**
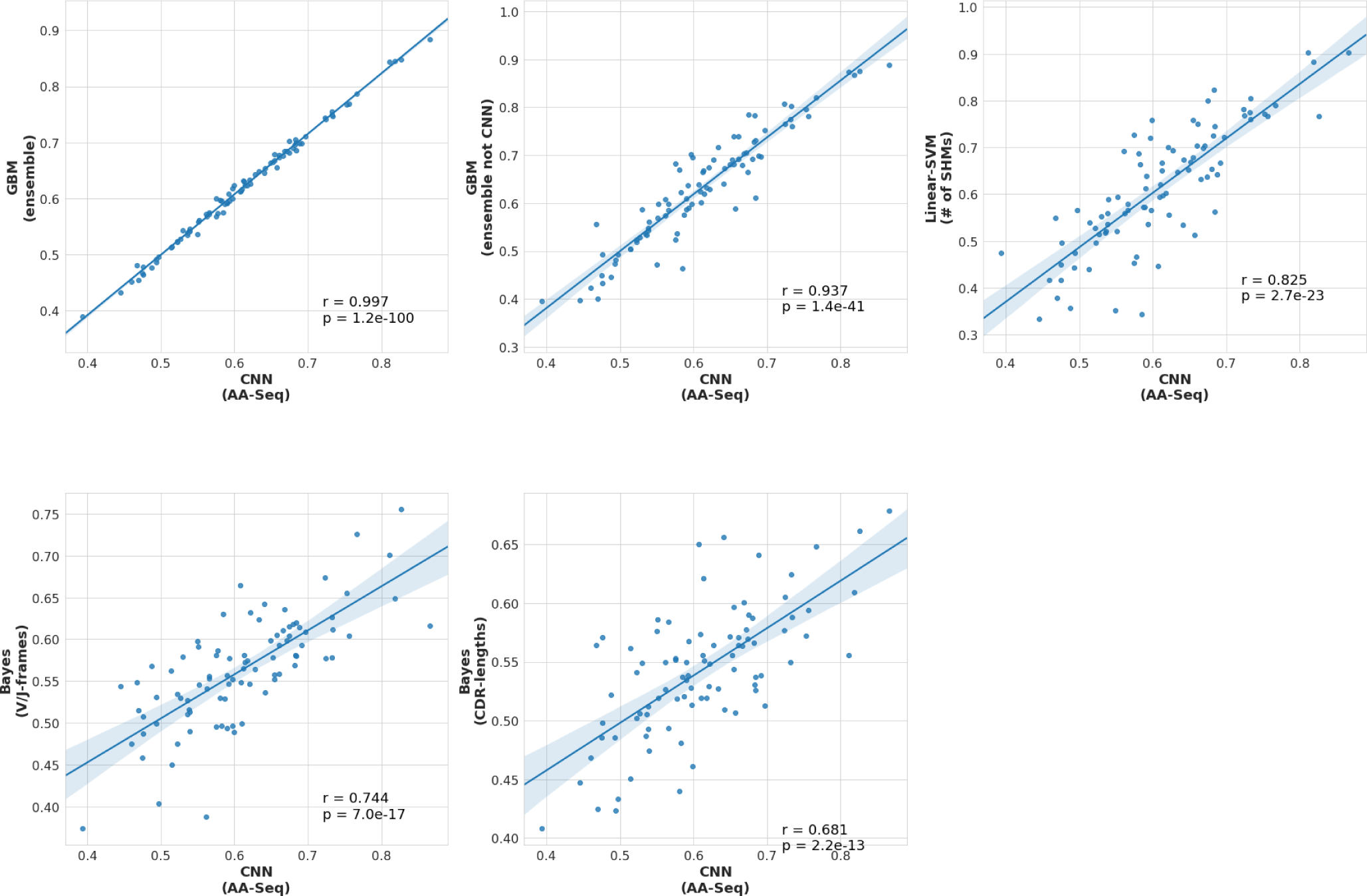
Scatter plots showing correlation between 89 AU-ROCs of CNN to those of other classifiers (ensemble classifier, ensemble classifier without CNN, linear classifier using # of SHMs, Bayes classifier using usage of V/J frames, and Bayes classifier using CDR-lengths.

Since input of CNN is the encoded physicochemical properties in amino acid sequences, and the classifiers is nonlinear, there is no straightforward way to extract biologically interpretable information from the model. Instead, sequence motifs in CDRs captured by the classifier were investigated, which is easy-to-understand. Sequence motifs on BCRs/Igs that were correctly classified or misclassified with high confidence in CDR1, CDR2, and CDR3 are shown in Figure 6. Unfortunately, no prominent motif patterns were found between the BCRs/Igs of normal and tumor tissues. Although there might be some differences between correctly classified motifs and misclassified ones, the biological importance is unclear.

The distributions of tumor probabilities in normal/tumor samples were investigated over all patients(Figure 5).

**Figure 5.**
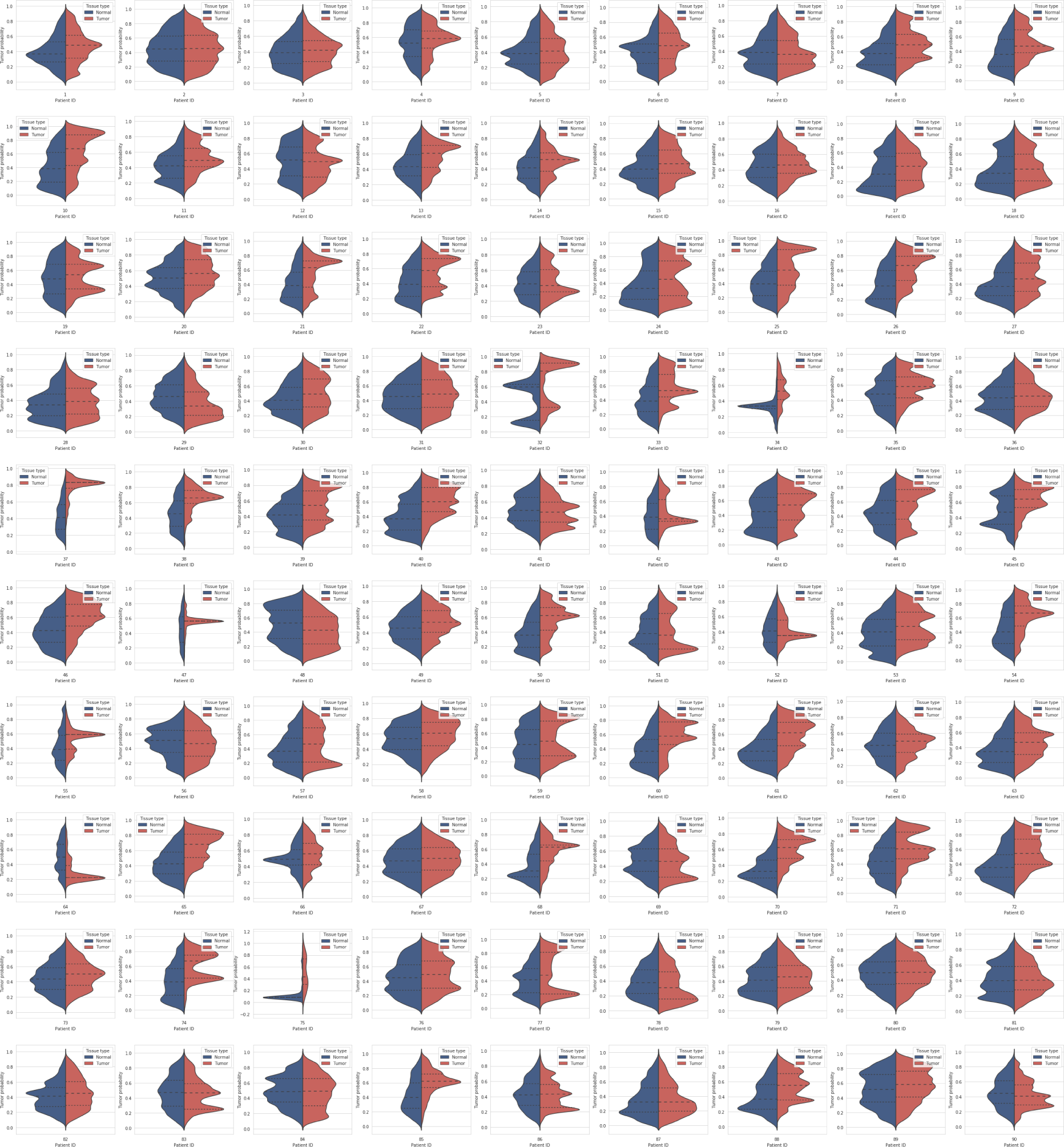
Clonal entropy and distribution of tumor probabilities in whole normal/tumor samples over 90 patients.

**Figure 6.**
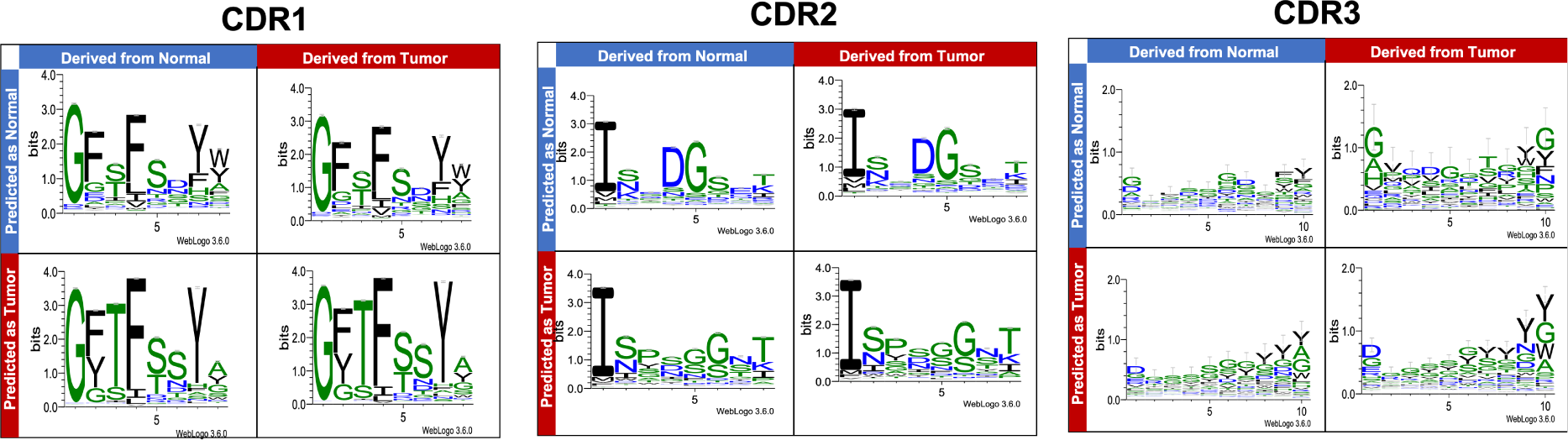
Sequence motifs constructed for all CDRs. Each motif is made using sequences that have average length of each region. We note that the y-axes of CDR3 is different from those of CDR1, CDR2.

**Figure 7.**
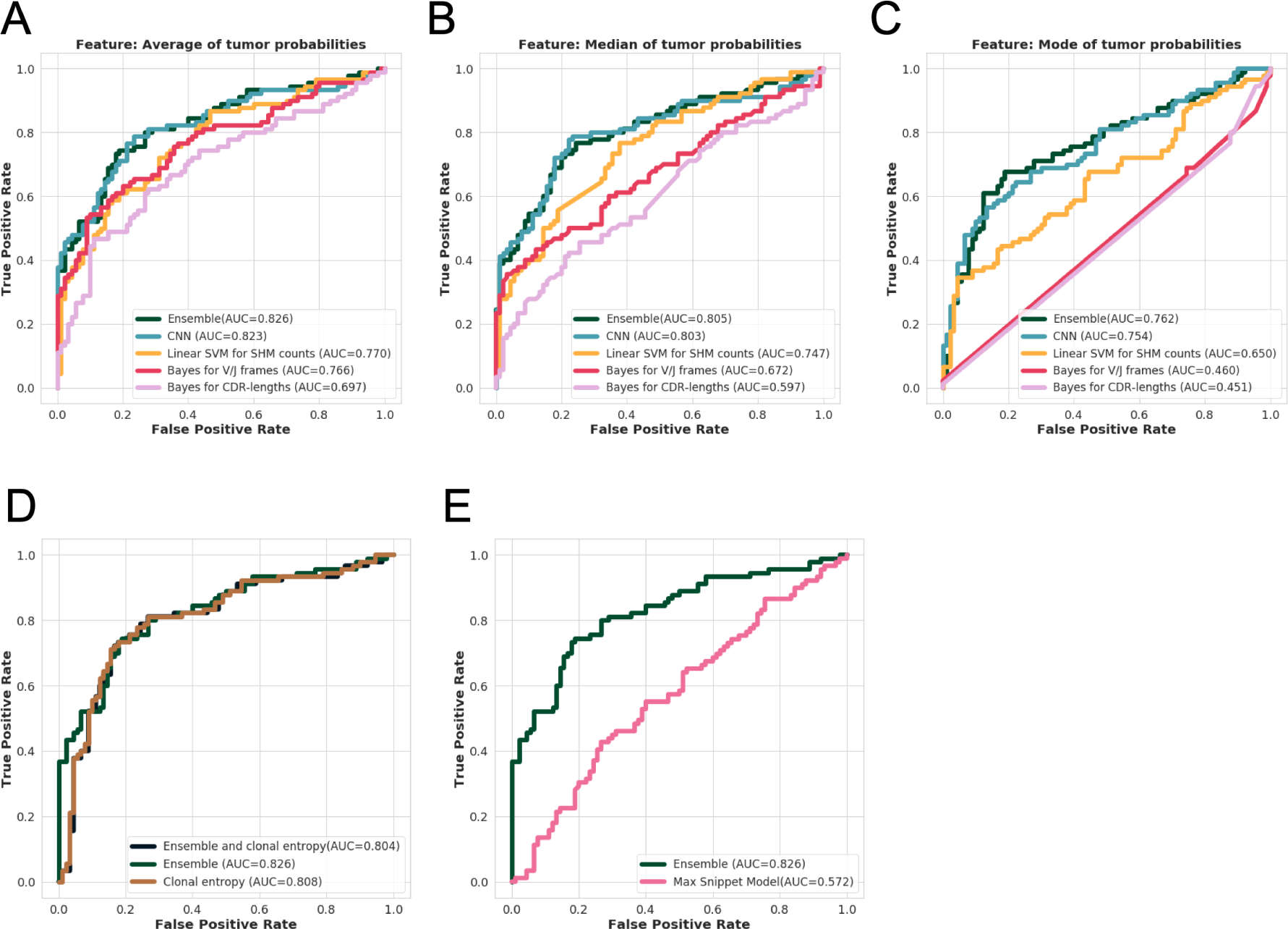
Receiver Operating Characteristic (ROC) curves showing tissue-level classification performance. Each figure illustrates the comparison of ROC-curves of (A) models using average score from sequence-independent features, (B) those using median score, (C) those using mode score, (D) ensemble model and clonal entropy, and (E) ensemble model and Ostmeyer’s model.

### 3.2 Tissue-level classification

Another main goal of this work was to precisely estimate tissue type, normal or tumor, using BCR/Ig sequences obtained from a tissue sample, under a hypothetical diagnostic scenario in which we can only access to a set of BCR/Ig sequences obtained from either of the tissue.

The tissue classifier was constructed using descriptive statistics values of the tumor probabilities of the already-trained GBM for individual BCRs/Igs. Figure 4A-C show the ROC-curves using average, median, and mode of the probabilities emitted by each single sequence-level classification models, respectively. Among the models based on the same sequence-level classifiers, the models employing average probabilities always outperformed the other two. Therefore, we adopted average probabilities as the descriptive statistics value for the tissue-level classification model in the rest of the study.

Tissue-level classification was also conducted by adding clonal entropy information to the ensemble model. However, the comparison of such models shown in Figure 4D suggested that it would not improve the classification performance.

In addition, our tissue-level classifier was compared to the previously introduced machine-learning method for disease classification based on BCR/Ig sequences^14^. Their model employs k-mer snippet on CDR3 sequences as a feature, and tissue-level classifications are carried out by constructing logistic regression model for a single snippet and by taking the maximum to aggregate the probabilities. Although this method was originally developed for classifying autoimmune diseases such as multiple sclerosis, it is applicable to cancer diagnosis. As shown in Figure 4E, our model significantly outperformed the Ostmeyer’s model in our cancer dataset (*p* = 3 × 10^−6^).

## 4 Discussion

In this paper, we hypothesized that BCR/Igs repertoire in tumor microenvironment have distinct features from those in normal microenvironment, and thus they can be distinguished using machine learning techniques from repertoire sequence data. Classifiers using various sequence-independent features were developed and compared the performances to investigate what features contribute to the difference of the repertoire.

In the first experiment, individual BCRs/Igs were classified using amino acid sequences of CDRs, lengths of CDRs, V/J-frames and the number of SHMs. We found that all the features contributed to the classification performance with different degrees. This result indicates that sequence-independent features of normal/tumor tissues have shared propensity across patients. We also found that the number of SHMs in tumor tissue is fewer than that in normal tissue and it is one of the most significant discriminative features. There are some possibilities which explain the observation. It might be because relatively short time has passed since immune system recognize the tumor antigens compared to other antigens in gastric tissues. Another possibility is that there is a mechanism that suppresses affinity maturation in tumor microenvironment. Although some of the mechanism are already known^26, 27^, the contribution in tumor immunity is yet to be investigated.

We also found that the CNN model using encoded amino acid sequences in BCRs/Igs contributed the most in the GBM ensemble model. High correlation value between the AUC values of the ensemble classifier without CNN model and that with CNN model suggests that CNN model captures most of the discriminative information captured by the other sequence-independent features. Although the performance gain over the classifier without CNN model was very small, the CNN model might be worthy of further investigation since there is still room for improvement. For example, our current CNN models cannot handle variable CDR lengths. Removing center of CDRs might lose important information for the discrimination. Incorporating more sophisticated machine learning models that can handle the variable length inputs such as Recurrent Neural Network might capture the information not capture by the other sequence-independent features and improve the classification performance.

In the second experiment, the tissue-level classification model was developed and compared the performance with Ostmeyer’s model, a logistic regression model using 6-mer snippets of CDR3 amino acid sequences in multiple-instance learning settings. Our model significantly outperformed Ostmeyer’s model. One of the advantages of our method is that it partially retains information about the position of amino acids in BCR/Ig sequences. Another advantage is that our model deals with global statistics of sequence-independent features, such as the number of SHMs and V/J-frame patterns. On the other hand, Ostmeyer’s model utilizes short sequence motifs in CDR3 sequences, irrespective of its relative position, and classification is performed based on only the most discriminative motif. The significant difference in the classification performance suggests that global sequence-independent features rather than individual amino acid sequences of BCRs/Igs contribute the difference between normal and tumor microenvironment, which could differ from other immune-related diseases such as autoimmune diseases. In autoimmune diseases, small number of common antigens and antibodies to the antigens contribute to the pathogenesis, which could result in the convergent amino acid sequence motifs in BCRs/Igs. On the other hand, in cancer microenvironment, antigens could be highly diverse both within each cancer tissue and across different cancer tissues. Such differences of immune landscapes can also explain why remarkable sequences motifs specific to normal/tumor microenvironments were not found.

The tissue-level classification in the second experiment achieved an AUC value of 0.826, which indicates that our model can be utilized in clinical applications. For example, cancer could be diagnosed even if the tissues do not contain cancer cells, so long as they contain tumor-infiltrated B cells or plasma cells. Indeed, single biopsies of gastric cancer tissues fail to show malignancy in 20% to 30% of cancer cases^28, 29^. Therefore, our approach could help correctly identify some of the false negatives in gastric biopsies and diagnose gastric cancers correctly by examining the BCR/Ig repertoires of B-cells adjacent to cancer cells, although further experimental validation will be required for clinical application of this concept.

## 5 Conclusions

Using a machine learning approach, we have shown that the BCR/Ig repertoire in gastric cancer tissues has distinct sequence characteristics compared to those of normal gastric tissues. BCR/Ig-features obtained in Rep-Seq showed their distinctiveness between normal and tumor environment in single sequence resolution and also showed high diagnostic capacity. Although we focused specifically on BCRs/Igs in gastric cancer, the same approach could be applied to evaluating TCRs and BCRs/Igs in other types of cancers or immune-related diseases, such as autoimmune diseases and infections, including those examined by past research^3–5, 7, 14^. Our analysis will open the door for new areas of tumor immunology and may lead to the development of novel diagnostic tools for cancer using repertoire sequencing.

